# Quantitative modeling of the function of kinetically driven transcriptional riboswitches

**DOI:** 10.1101/821884

**Authors:** César Parra-Rojas, Boris Fürtig, Harald Schwalbe, Esteban A. Hernandez-Vargas

## Abstract

We propose a generalized modeling framework for the kinetic mechanisms of transcriptional riboswitches. The formalism accommodates time-dependent transcription rates and changes of metabolite concentration and permits incorporation of variations in transcription rate depending on transcript length. We derive explicit analytical expressions for the fraction of transcripts that determine repression or activation of gene expression, pause site location and its slowing down of transcription for the case of the (2’dG)-sensing riboswitch from *Mesoplasma florum*. Our modeling challenges the current view on the exclusive importance of metabolite binding to transcripts containing only the aptamer domain. Numerical simulations of transcription proceeding in a continuous manner under time-dependent changes of metabolite concentration further suggest that rapid modulations in concentration result in a reduced dynamic range for riboswitch function regardless of transcription rate, while a combination of slow modulations and small transcription rates ensures a wide range of finely tuneable regulatory outcomes.

## 1. Introduction

Riboswitches are non-coding RNA sequences that up- or down-regulate gene expression by selectively binding metabolites and subsequent conformational transitions without the intervention of protein factors [1]. They commonly reside in the 5’-untranslated region of bacterial mRNAs. Typically, riboswitches contain two functional domains: an *aptamer domain*, which forms the metabolite-binding pocket, and an *expression platform*, which interacts with the aptamer, assessing the metabolite binding status and modulating gene expression via conformational changes in response to it [1, 2]—see Fig. 1.

**Figure 1:**
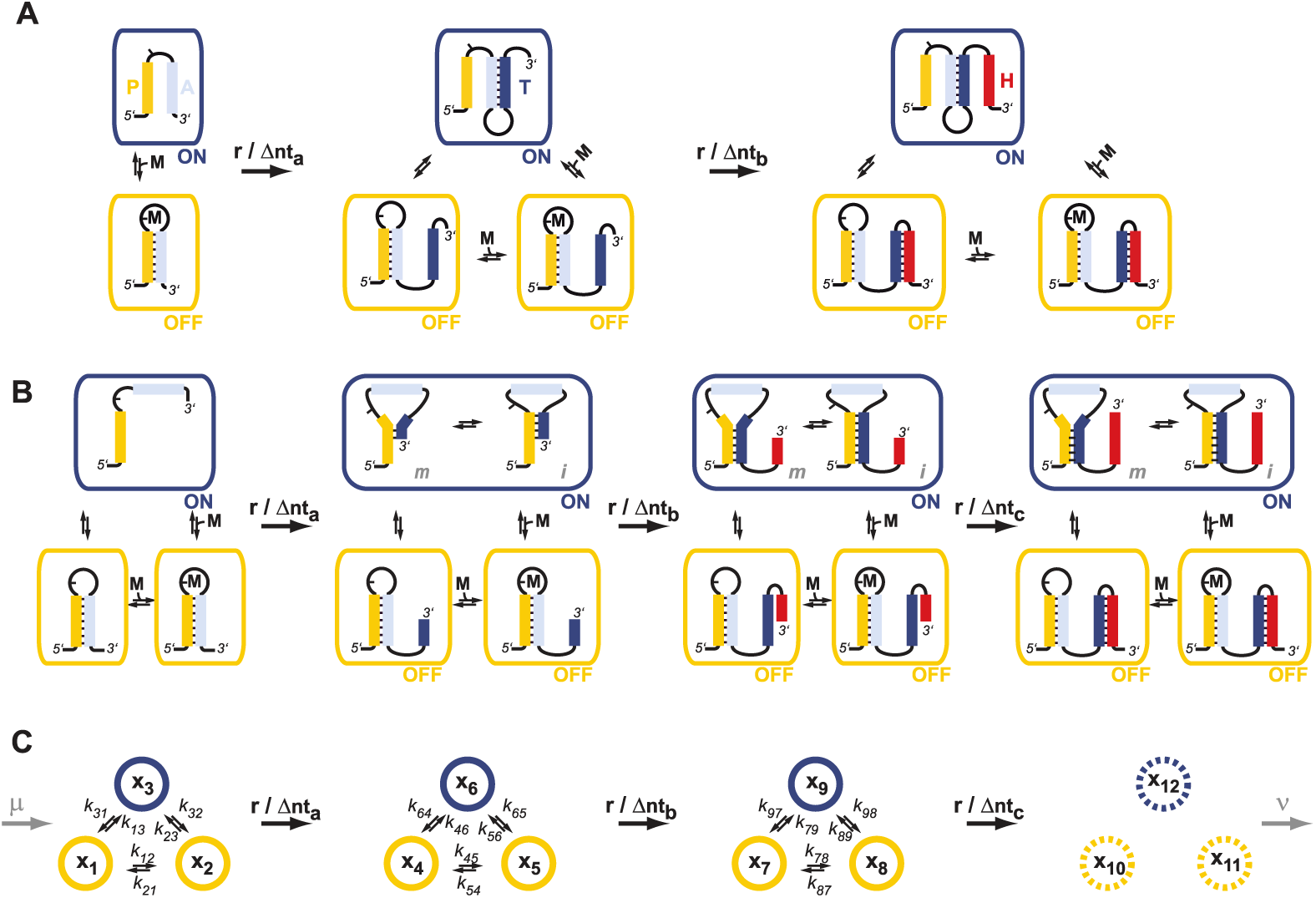
Models of co-transcriptional folding pathways **(A)** Conformational states in context of the transcription progress of a riboswitch with a sequential base-paired stabilized arrangement of the switching sequences P, A,T and H (yellow, grey, blue and red, respectively; arrangement is exemplified by the guanine sensing riboswitch). Metabolite (M, black) binding can occur as soon as the aptamer domain is transcribed and consequently locks A in the PA helix resulting in the population of a single conformation during the transcription process and subsequently in transcription termination. The aptamer domain is synthesized first. As transcription continues, the free mRNA adopts the metastable antiterminator conformation stabilized by AT interaction which refolds to the terminator conformation after the riboswitch is completed. **(B)** Conformational states in context of the transcription progress of a riboswitch with a non-sequential base-pair stabilized arrangement of the switching sequences P, A, T and H (exemplied by the deoxy-guanosine sensing riboswitch). Upon transcription of sequence element T the riboswitch can adopt two fastly interconverting conformations (m and i) in the metabolite free functional ON-state. For both types of riboswitches the functional OFF-state is always characterized by stable interactions between P and A, whereas all functional OFF-states favour non base-paired H sequences. **(C)** The abstraction for the mathematical model, where the m and i ON states are merged into one, so that *N*_*on*_ = 1 and *N*_*off*_ = 2. The number of intermediates is *L* = 3, and sink states are introduced at the regulatory decision point, totalling *N* = 12 states for the model. The transcription rate, in nucleotides per second, is denoted by *r*, while the length of the separation between intermediates, in nucleotides, is denoted by Δnt_*a,b,c*_; at the same time, *µ* and *ν* can be thought of as the rates of transcription initiation and transcript degradation, respectively, and are introduced in the model when continuous transcription is considered.

Riboswitches can act at the level of transcription or translation. Their regulatory function is achieved mainly by two mechanisms corresponding to translation initiation and rho-independent transcription termination or antitermination. In the case of translation, regulation involves the ribosome binding site (RBS) of the mRNA, which can be either shielded from (OFF switch) or accessible to (ON switch) ribosome binding to the mRNA [3] in response to metabolite binding to the upstream aptamer domain. Repression of transcription involves the conditional formation of a terminator stem in response to metabolite binding. This results in premature abortion of transcription. If the terminator is not formed by facilitating of a mutually-exclusive *antiterminator* structure [1, 4], RNA transcription continues. The interaction between the two functional domains, the aptamer domain and the expression platform, relies on the base pairing behavior of generally four complementary nucleotide stretches: P, A, T, and H. In the OFF state, the 5’-aptamer-strand (P) pairs with the aptamer-stabilizing strand (A) and the switching strand (T) forms the terminator helix together with the terminator strand (H). By contrast, in the ON state, A and T or P and T pair and form the antiterminator conformation, and the P/A and H strands are unpaired.

Regulation of transcription by riboswitches is a kinetic phenomenon that cannot be explained by assuming only a few regulatory relevant static riboswitch conformations and the differences of binding affinities at equilibrium to these states. The slow kinetics of RNA refolding after ligand binding play an essential role in the process [5]: if binding does not occur before the terminator stem is synthetized, transcription will proceed. In biological assays, metabolite concentrations 2-3 orders of magnitude higher than the equilibrium dissociation constant *K*_*D*_ are required to effectively regulate expression and, the faster the transcription, the higher the concentration needed in order to trigger termination [6]. Such high affinities of riboswitches for their corresponding metabolites can be exceedingly high, such that an aptamer could potentially detect even a single molecule per cell [4]. This would render gene expression regulation infeasible. Therefore, manipulating the transcription speed by means of *e.g.*, lengthening or shortening the time needed for the formation of the terminator stem conformation [4], or by exploiting pause sites [2], results in tuning the concentration levels at which the riboswitch is able to operate. Incorporation and location of pause sites for transcriptional regulation underlies strict requirements: They have to be placed at specific positions within the transcript, as they have to modulate the time frame for synthesis intervening sequences A-to-T and T-to-H.

The transcription rate not only competes against metabolite binding rates, but also against the refolding rates of RNA itself, and the effects of transcription speed on gene expression can be observed even in total absence of metabolite [7]. These general considerations highlight the importance of the formation of metastable conformations during transcription, and the existence of a well-defined range of transcription speeds at which riboswitch function is possible at all [8]. The simplicity of the regulatory function of riboswitches and the accessibility of all required kinetics and thermodynamics for such regulation at least *in vitro* allows for quantitative modeling of regulation function. Here, we develop a model that integrates all regulatory important factors: transcription speed and its modulation by pause sites, acquisition of RNA structure, metabolite-binding-induced RNA refolding, with their associated timescales, as well as the influence of metabolite concentration, and external variables such as temperature. While the analysis shown here is based on the particular example of an OFF switch, the modeling formalism is applicable to both ON and OFF transcriptional riboswitches in general.

## 2. Materials and Methods

### 2.1. Riboswitch folding kinetics

We consider a system of *L* different transcriptional intermediates, with *N*_*on*_ ON states and *N*_*off*_ OFF states at each of these steps, the latter including both ligand-free and ligand-bound states—we base the description on an OFF switch, and discuss the ON case below. We will be interested in the fraction of mRNA chains that remain in the ON state during transcription, *i.e.*, the fraction ON among the full-length transcripts. In practice, this means that we need to allow for complete conversion of the shorter transcription intermediates. We introduce *sink* states at the end of a single round of transcription, that will accumulate the transcripts in ON and OFF states after the regulatory decision point has been reached and transitions between states are no longer possible or no longer biologically relevant in terms of gene expression. In mathematical terms, the quantity of interest corresponds to the combined steady-state values of the ON sinks. The total number of states in the system will correspond to *N* = *N*_*L*_(*L* + 1), where *N*_*L*_ = *N*_*on*_ + *N*_*off*_ is the total number of states per transcript length. The fraction of transcripts occupying state *i* at a given time *t* will be denoted by *x*_*i*_(*t*), so that the occupation levels in the model are characterized by a vector ***x***(*t*) = (*x*_1_(*t*), …, *x*_*N*_ (*t*)). An illustration of such a system with *L* = 3, *N*_*on*_ = 1, and *N*_*off*_ = 2—thus, *N* = 12—is shown in Fig. 1. An important distinction must be made between this formulation and our previous works [9, 7, 8], which do not consider a final set of states and therefore were simulated up to a *stopping time* given by the mean elongation time from the shortest to the longest transcripts.

The general form of the kinetics is given by

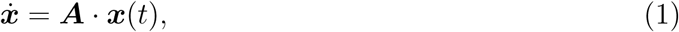

where ***A*** is an *N* × *N* matrix with entries

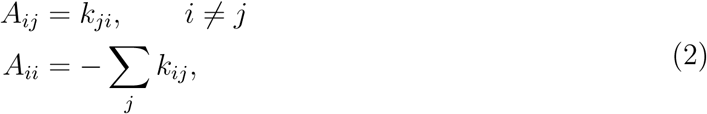

and *k*_*ij*_ represents the folding rate from state *i* to state *j*. Here, we assume that the folding rates between conformations have been either measured experimentally or predicted *in silico*—see, *e.g.*, Flamm *et al.* (2002) [10]. The number of relevant intermediates, *L*, and the distance in nucleotides between any given pair of them, is chosen *a posteriori* so that the *k*_*ij*_ are all constant. This means that, *e.g.*, in Fig. 1 the folding rates were measured and found to remain constant or approximately constant for Δnt_*a*_ nucleotides, then changing and remaining the same for another Δnt_*b*_ nucleotides, and so on.

We will adopt the convention that states are ordered from short to long, from OFF to ON so that, for a system of size *N*, we will have

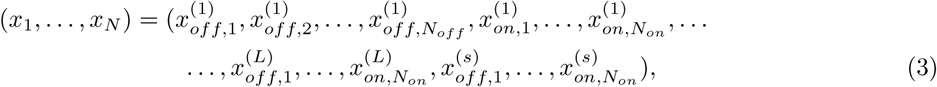

where *s* stands for ‘sink’. We arrange the state vector in the mathematical description so that, per transcript intermediate, all ligand-free states come before ligand-bound states. From this, it follows that the quantity of interest will be the last *N*_*on*_ elements of the state vector when *t* → ∞. If we decompose the label of a certain state *i* as *i* = *p*_*i*_*N*_*L*_ + *q*_*i*_, with *p*_*i*_ = 0, 1, …, *L* and *q*_*i*_ = 1, 2, …, *N*_*L*_, then the following properties are satisfied:

1. All transitions rates *k*_*ij*_ for which *p*_*i*_ > *p*_*j*_ are equal to zero; that is, transitions from longer to shorter transcripts are forbidden.
2. Transitions for which *p*_*i*_ < *p*_*j*_ and *q*_*i*_ ≠ *q*_*j*_ are also equal to zero, since the lengthening of the transcripts preserves their state. In other words, the lengthening of a transcript in, *e.g.*, state 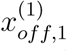 can only take it to state 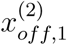 but not to, say, 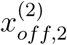
3. Transitions for which *q*_*i*_ = *q*_*j*_ but *p*_*j*_−*p*_*i*_ > 1 are also equal to zero, due to the fact that the lengthening is sequential.
4. By construction, the transition rates between two given transcript lengths are all equal and independent of the state, so that *k*_*ij*_ = *k*_*lm*_ for all cases when *p*_*i*_ = *p*_*l*_ and *p*_*j*_ = *p*_*m*_. Furthermore, these rates are uniquely determined by the transcription rate, which we denote by *r*; *r* has units of nt · s^−1^, and is defined as *r* = 1 [nt] · *K* [s^−1^], where *K* represents the rate of nucleotide addition by the polymerase, in s^−1^.

### 2.1.1. Analytical solutions

The model can be explored numerically, both for constant and time-varying transcription rates. However, analytical solutions can be found if the transcription rate is a constant and the total number of states is sufficiently small. The general solution reads

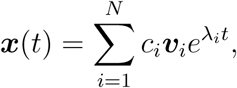

where λ_*i*_ and ***v***_*i*_ are, respectively, the *i*-th eigenvalue and the *i*-th right eigenvector of ***A***, while the *c*_*i*_, *i* = 1, …, *N* are coefficients that are uniquely determined by the initial condition ***x***_0_ = ***x***(*t* = 0). Using the expression above, these can be found from

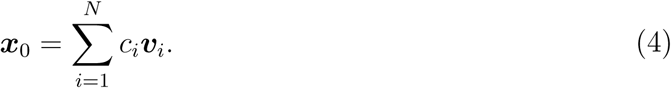

We note that, by construction, the system with the sink states will have *N*_*L*_ eigenvalues equal to zero with eigenvectors being the unit vectors corresponding to the sinks—*v*_*i,j*_ = *δ*_*ij*_ for *i* = *N*_*L*_*L* + 1, …, *N* —and with all other eigenvalues λ_*i*_ < 0. Thus, its steady-state solution, which we denote by a vector 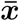, will be given by

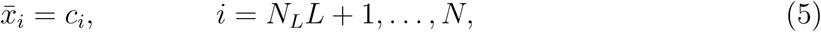

with 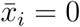 otherwise.

Let us consider the (2’dG)-sensing riboswitch from *Mesoplasma florum* as an illustration. Previously, we have determined experimentally the relevant transcriptional intermediates and the folding rates between conformations for this riboswitch [8]. In principle, this system consists of two ON states per transcriptional intermediate, with *L* = 2 transcriptional intermediates; however, if we are interested in the total fraction of transcripts allowing gene expression, the system size effectively reduces from twelve to nine dimensions. This is analogous to Fig. 1B, showing two ON states merged into one for a system with *L* = 3, reduced from 16 to 12 states. As in the original model, let us assume that there is a binding window of length Δnt_*a*_ ≡ 1*/a* that has already been traversed, with a resulting fraction *b*_0_ of bound transcripts. That is, we consider only the states *x*_4_ to *x*_9_ of the nine-dimensional representation, with an initial condition given by: *x*_4_ = 1 *− b*_0_, *x*_5_ = *b*_0_, *x*_*i*_ = 0 for *i* = 6, …, 9. This corresponds to a single round of transcription starting from its shortest length—*x*_*i*_ = 0 for *i* = 7, 8, 9—and in a fully OFF configuration—*x*_6_ = 0—with a fraction *b*_0_ of bound transcripts. Folding is only possible from OFF to ON states—the reverse rates are negligible [8]—so that we set *k*_45_ = *k*_54_ = *k*_64_ = *k*_65_ = 0, and an explicit solution can be found for the fraction ON at full length:

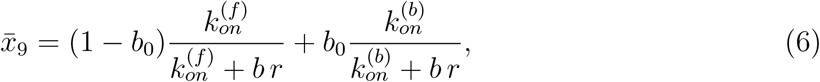

where Δnt_*b*_ ≡ 1*/b* = 24 nt is the length of the folding window from nucleotide 113 to nucleotide 137, 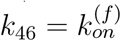 is the folding rate from the ligand-free OFF state to the ON state, and 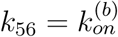 is the folding rate from the ligand-bound OFF state to the ON state. We may compare this result to the original [8], given by

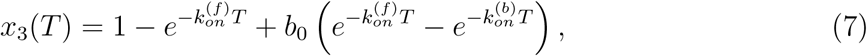

with *T* = (*b r*)^−1^ s, corresponding to the mean time it takes the transcription to proceed by 24nt.

### 2.2. Pause sites

We start by noting that changes in the stability of the different conformations enter the model through the addition or removal of intermediates of a given length. In other words, the intermediates are defined by windows of approximately constant folding rates so that, by construction, all rates *k*_*ij*_ appearing in ***A***—Eq. (2)—and relating to transitions between conformations are constant, while elongation transitions are directly defined through the transcription rate *r*.

For the model systems described here, we have previously shown that the transcription rate experiences changes at specific transcript lengths. In this case we can exploit the same strategy as for changes in folding rates: we can insert a window of reduced transcription rate as another transcriptional intermediate in the system. In other words, in order to model a pause site located, *e.g*, in the last quarter of a window of length Δnt, we can insert an intermediate that divides the window into one sub-window of length 3Δnt*/*4 and another of Δnt*/*4, with the latter being transcribed at a (much) lower rate. In general, changes in transcription rate at specific locations can be accommodated by further subdivisions.

### 2.3. Metabolite concentration changes

Another factor that influences the dynamics of the system is the metabolite concentration, which will have an impact on the fraction of bound transcripts at the end of the binding window, and that we have so far not considered. Since the metabolite does not play a role in the fate of the transcripts once they are past the binding window, in order to explore the consequences of varying its concentration over slower timescales we need to modify the model to account for continuous transcription. We can picture this as a system that is producing transcripts at a rate *µ*, that we may identify with the rate of *transcription initiation*; at the same time, we assume that full-length transcripts leave the system at a rate *ν*, signifying *transcript degradation*—see Fig. 1. We make a simplification by treating the metabolite concentration *M* as an external reservoir we can modify arbitrarily; that is, it does not depend on the dynamics of the riboswitch. Additionally, for the sake of simplicity, we take the degradation rate to be the same as the initiation rate, *ν* = *µ*. Under this condition, the total sum of full-length transcripts reaches unity at equilibrium, and we will be interested in the fraction of full-length transcripts in ON state once this equilibrium has been reached.

We initialize the system in absence of metabolite, and start monotonically increasing its concentration at a certain time *t*_0_ up to a time *t*_1_, such that if *t*_1_ >> *t*_0_ the concentration reaches a saturation value which we choose to be *M* = 1. At time *t*_1_, we begin to decrease the concentration until no metabolite is present in the system. We use

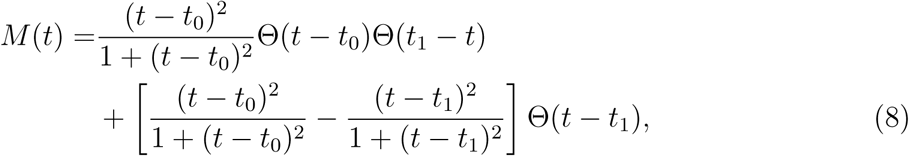

where Θ (·) corresponds to the Heaviside theta function.

## 3. Results

### 3.1. *Full conversion of shorter transcripts in the (2’dG)-sensing riboswitch from* Mesoplasma florum *results in a lower fraction ON*

The analytical solution for the fraction ON at a length of 137 nucleotides in the (2’dG)-sensing riboswitch from *Mesoplasma florum* is shown in Eqs. 6 and 7 for the formulations with and without sink states, respectively. Comparing both results, we observe a generally lower fraction ON in the case with sinks—see Fig. 2. We can explain this discrepancy by noting that, in the original formulation [8], all transcripts at nucleotide 113 have a life-time that is exactly equal to the mean time to elongation by 24nt, *T*. By contrast, in the case with sinks, the life-times are exponentially distributed with the same mean. As a result, the majority of the transcripts will be elongated before a time *T* has passed, and have therefore a lower chance of folding towards the ON state. This is a subtle but important conceptual difference between this and our previous works [9, 7, 8].

**Figure 2:**
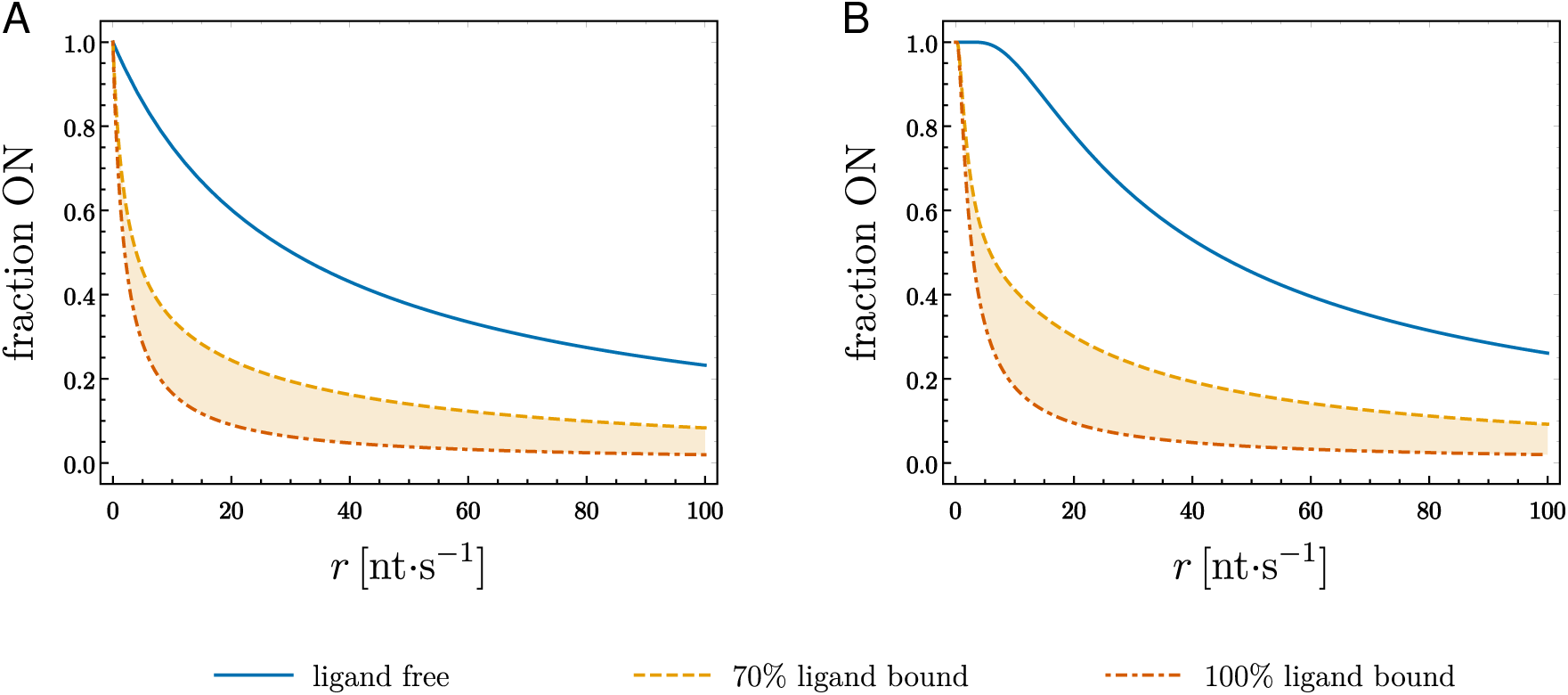
Fraction of transcripts in the ON conformation at a length of 137 nucleotides in a single round of transcription for the (2’dG)-sensing riboswitch from *Mesoplasma florum*, as a function of constant transcription rate *r* and a fixed fraction of initially bound transcripts. **(A)** With full conversion of shorter transcripts through the addition of sink states. **(B)** In the original formulation without the sink states, from Helmling *et al.* (2018) [8].

We have started from a specified fraction of ligand-bound transcripts for the purpose of the comparison. However, the full result must take into account the fact that this fraction is the result of binding events between the first and second transcriptional intermediates— see Fig. 1. Therefore, we include the dynamics during the binding window, with allowed transitions only from the unbound to the bound OFF state—*k*_13_ = *k*_21_ = *k*_23_ = *k*_31_ = *k*_32_ = 0 in Fig. 1. We take *k*_12_ = *k*_*b*_ *M*, where *k*_*b*_ is the metabolite binding rate and we have assumed that the ligand concentration, *M*, is large enough so that binding events have a negligible effect on it. In this case, 1*/a* = 35 nt, and 1*/b* = 24 nt, corresponding to the length of the binding and folding windows, respectively. The full solution for 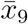, corresponding to the fraction ON for a transcript length of 137 nucleotides in a single round of transcription— *µ* = *ν* = 0—takes the form

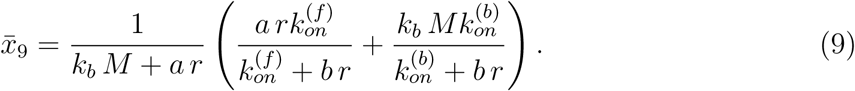

The full landscape of this solution is plotted in Fig. 3A as a function of a constant transcription rate *r* and the product *k*_*b*_ *M*. It is straightforward to see that the limits *k*_*b*_ *M* → 0 and *k*_*b*_ *M* → ∞ result, respectively, in no ligand binding and 100% ligand binding, corresponding to the cases *b*_0_ = 0 and *b*_0_ = 1 from Eq. 6. For vanishingly slow transition rates, the resulting fraction ON tends to unity regardless of the ligand concentration and binding rate.

**Figure 3:**
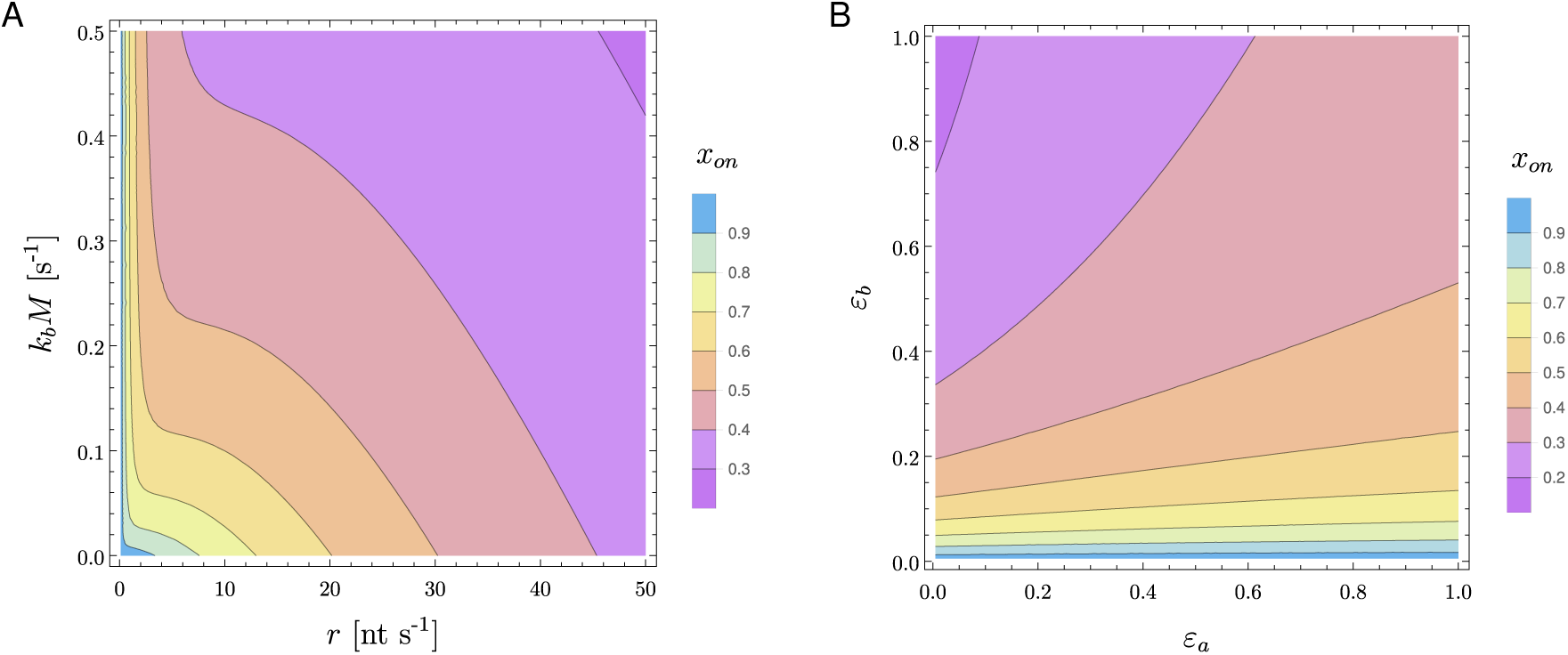
Contour lines for the fraction ON at a length of 137 nucleotides in a single round of transcription for the (2’dG)-sensing riboswitch from *Mesoplasma florum*—same as Fig. 1B, but with *L* = 2, *N* = 9—with folding rates taken from Helmling *et al.* (2018) [8]. **(A)** As a function of constant transcription rate *r* and the product of the ligand binding rate and the ligand concentration *k*_*b*_ *M*, obtained from Eq. 9. **(B)** With binding (length 1*/a*) and folding (length 1*/b*) windows divided into two sections traversed at different transcription rates, 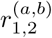. Result obtained from Eq. 11 when the rates follow 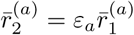, and 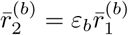, and are expressed as a function as pause site location and its amount of slowing down: respectively, either at the binding (*ε*_*a*_ < 1) or folding (*ε*_*b*_ < 1) window, and the exact magnitude of *ε*_*a,b*_. Base parameters are 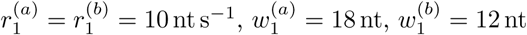, and *k*_*b*_ *M* = 10*/*18 s^−1^.

Something the results show is that solutions can be found by iteration. Suppose that we focus only on the binding window of length 1*/a*, and “stop” the dynamics after it is traversed. If we identify *b*_0_ as the solution for *x*_5_ when *t* → ∞ if we forbid all transitions between *x*_4_, *x*_5_ and *x*_6_, we obtain

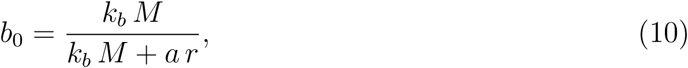

and Eq. 9 reduces to Eq. 6. In other words, a full analytical solution for intermediate states at a given length may be found recursively from the solution to the fraction of transcripts that eventually pass through the states at the previous length.

### 3.2. Pause sites located after the binding window increase the switching efficiency

As an illustration, both the binding and folding windows of the system from the previous section are divided into two sub-windows, the lengths of which we denote by *w*_1_ and *w*_2_; that is, 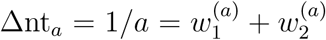, and 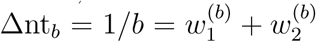. We assume that, in each case, the sub-window *w*_1_ is transcribed at rate *r*_1_ and the sub-window *w*_2_ is transcribed at rate *r*_2_. We are interested in the effects of the location of pausing events, which will enter as a difference between the rates *r*_1,2_ in the binding window, the folding window, or both. We are also interested in the intensity of the slowing down in transcription, which will correspond to how much lower is one of the rates *r*_1,2_ with respect to the other.

This modified system has a total of 15 states, and the steady-state solution for the final fraction ON in a single round of transcription is given by

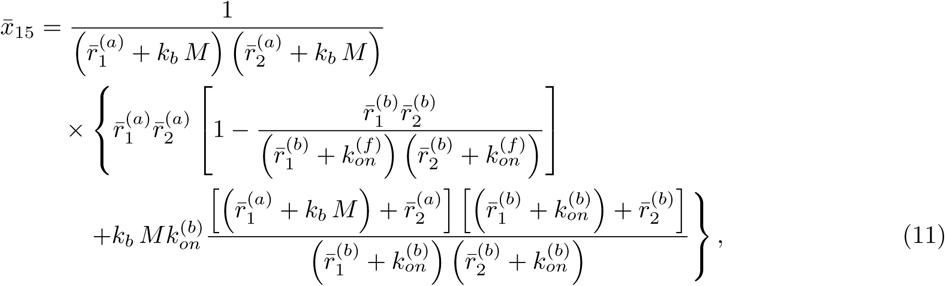

where 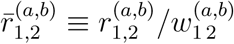. This solution reduces to Eq. 9 when all the transcription rates are equal, one of the binding windows takes the full length 1*/a*, and one of the folding windows takes the full length 1*/b*.

Since a more pronounced slowing down of transcription has the same effect as lengthening the pausing window—note that the terms involving the transcription rates, 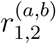, appear only in combination with those corresponding to window length, 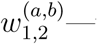we can explore the effects of pausing strenght and extension at the same time. Let us suppose that 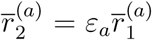, and 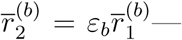that is, the effective transcription rate at one portion of the binding/folding window is a well-defined fraction of the one at the other portion. Fig. 3B shows the resulting fraction ON from Eq. 11 as a function of this fraction. We see from this that stronger pause events localized in the binding window tend to result in a lower fraction ON, due to the transcripts having more time to bind the metabolite. Stronger pauses during the folding window, in turn, have the opposite effect, since the transcripts have more time to refold from the OFF to the ON state. In agreement with the results depicted in Fig. 3A, a sufficiently intense reduction in transcription rate during folding results in the fraction ON being close to unity: 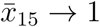 in Eq. 11 for *ε*_*b*_ → 0, regardless of the value of *ε*_*a*_.

We have explored the simple case of a pause extending from the beginning of the window until a certain point (resp. from a certain point until the end). The more realistic scenario of a short localized pause can be studied using the recursive property of the solutions that we have discussed above. However, we expect the qualitative insights gleaned from Eq. 11 to be valid in this case as well.

### 3.3. Fast variations in metabolite concentration reduce the dynamic range

As before, we take the folding rates from our previous work [8]. We further choose *k*_*b*_ = 1 so that, for a constant metabolite concentration *M* = 1 and a constant transcription speed *r* = 35 nt s^−1^, the fraction of bound transcripts at nucleotide 113 is *b*_0_ = 0.5 if initiation and degradation are ignored—see Eq. 10. As an example, we take the metabolite concentration to satisfy Eq. 8. Fig. 4 shows *M* (*t*), along with the resulting dynamics for the fraction ON at full-length for a choice of *t*_0_ and different values of *t*_1_ and transcription rate. In this illustration we see that, the larger the transcription rate, the faster the system tends to react to changes in metabolite concentration. However, at the same time, the folding window is traversed too quickly for conformational changes to occur to a great extent even for vanishing *M*, so that the dynamic range of the output is reduced as we increase the transcription rate. On the other hand, smaller transcription rates result in a slower reaction time but permit increased refolding. In this case, slower modulations in *M* allow the system to equilibrate at a larger fraction of bound transcripts at the end of the binding window. The combination of both larger concentration equilibration times and slower transcription rates results in a wider dynamic range for riboswitch function.

**Figure 4:**
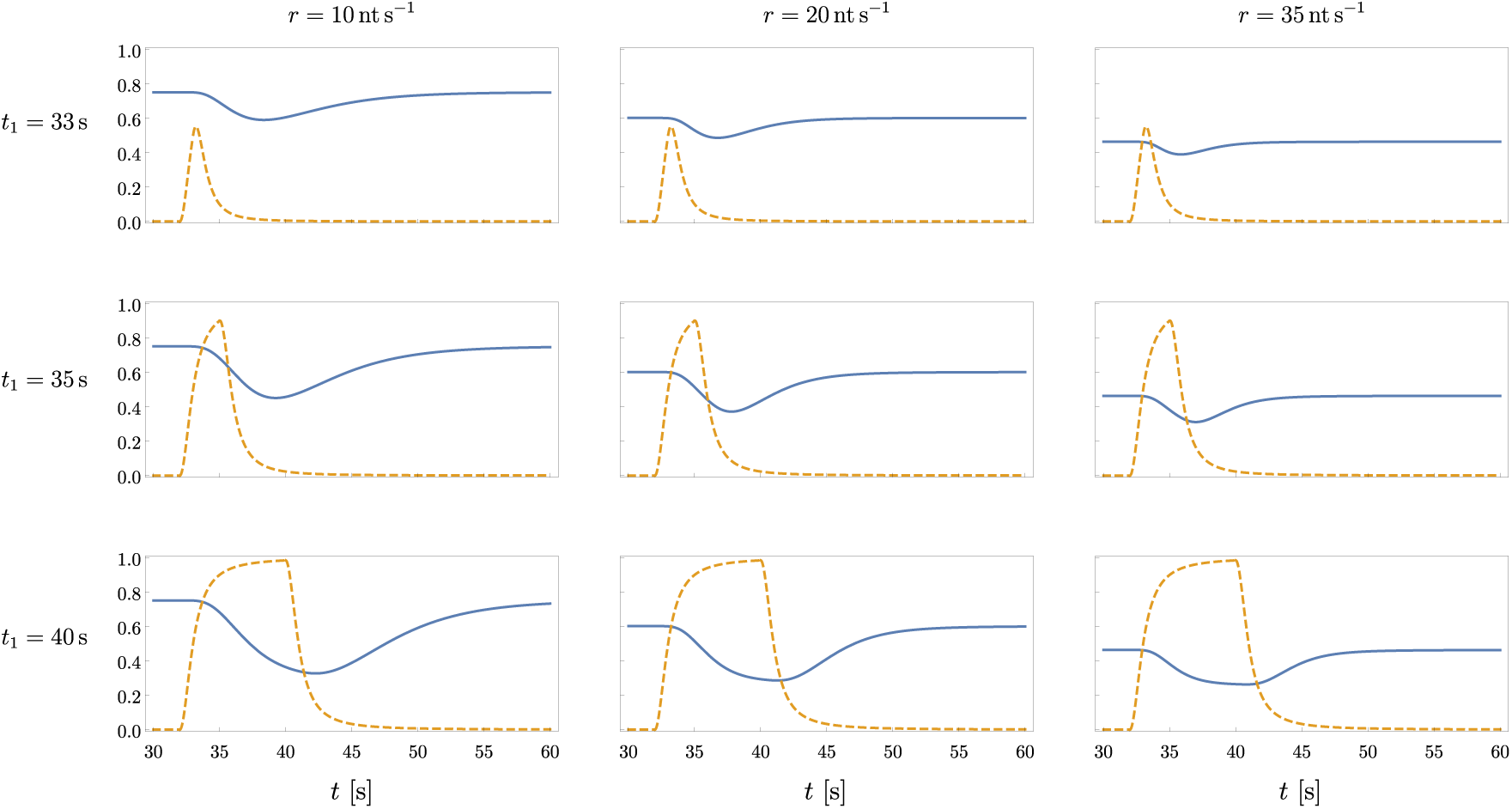
**Orange, dashed line:** time-varying ligand concentration from Eq. 8 for *t*_0_ = 32 s; top: *t*_1_ = 33 s; middle: *t*_1_ = 35 s; bottom: *t*_1_ = 40 s. **Blue, solid line:** resulting fraction ON at a length of 137 nucleotides for the (2’dG)-sensing riboswitch from *Mesoplasma florum* under continuous transcription at a constant rate; left: *r* = 10 nt s^−1^; center: *r* = 20 nt s^−1^; right: *r* = 35 nt s^−1^. An equilibration time of 30 s has been used. Parameter values: *k*_*b*_ = *µ* = *ν* = 1. Refolding rates from Helmling *et al.* (2018) [8].

## 4. Discussion

Naturally-occurring riboswitches are commonly found in bacteria, where they take part in the regulation of the expression of a number of genes which are essential for survival [11], as well as directly influencing motility and pathogenesis [12]. This, coupled with the selectivity with which they bind their corresponding metabolites, makes them attractive drug targets for the design of novel antibiotics [13, 11]. Engineered riboswitches, in turn, have been successfully used to control the replication and infectivity of both RNA and DNA viruses [14, 15]. This highlights a variety of potential therapeutic applications of riboswitch regulation, and the need for a deep qualitative and quantitative understanding of the mechanisms involved.

Mathematical approaches to riboswitch kinetics are scarce in the literature: examples include Santillán & Mackey (2005) [16], who constructed a dynamical model for the B_12_-sensing *btuB* riboswitch from *E. coli* and analyzed transient and steady states in terms of B_12_ levels subject to both transcriptional and translational regulation; Beisel & Smolke (2009) [17], who qualitatively outline the general performance properties, in terms of protein levels, of ON and OFF riboswitches under different regulatory mechanisms, and their implications for the tuning of natural and synthetic riboswitches; and Hofacker *et al.* (2010) [18], who describe the dynamics of RNA co-transcriptional folding in a step-by-step formulation through a set of transcription stopping times.

Here, we have presented a generalized modeling framework for the kinetics of transcriptional riboswitches, based on a group of relevant transcriptional intermediates defined by windows of approximately constant (re)folding rates, with a number of states at each step defined by the different available structural conformations. We assume that these windows are determined experimentally, and the relevant intermediates are subsequently chosen based on them. We introduce a final set of states, with no folding transitions between them, so that the output of the model corresponds to fraction of transcripts that eventually exit a transcription round in a given conformation. The formulation of the model is general and simple enough to accommodate either ON or OFF switches, and we note that, while alternatives for a full numerical characterization of co-transcriptional folding are available [19], our aim is to provide a framework to readily try out different configurations in a system with known, few relevant states, which permits both qualitative and quantitative analyses of the co-transcriptional interplay between transcription and folding rates, as well as metabolite concentration and binding rate. The sequential character of the model allows for a closed-form solution to be obtained at each intermediate step for sufficiently simple conformation configurations, *e.g.*, through the use of computer algebra [20], while the full solution can be subsequently pieced together from its component parts in a recursive manner. Furthermore, the existence of closed-form solutions facilitate the direct qualitative and quantitative exploration of the behavior of *classes* of riboswitches sharing their main structure and differing only on their distance between intermediates or their folding rates.

As an illustration, we have derived a number of analytical expressions for the fraction of transcripts in ON state at the regulatory decision point for a simple system, and applied them to the explicit case of the (2’dG)-sensing riboswitch from *Mesoplasma florum*; the model is able to reproduce previous results when the condition of short-transcript conversion is relaxed [8]. Due to the recursive character of the solutions, we have also shown how changes in conformation stability and position-dependent changes in transcription rate can be incorporated in the model through additional intermediates without rendering the system intractable. With this, we derived an analytical expression for the landscape of possible outcomes representing the effects of pause site strength and location in the system. For the case of the (2’dG)-sensing riboswitch from *Mesoplasma florum*, pausing at latter stages in transcription has a much stronger effect on the outcome than pausing during the binding window: while metabolite concentration and binding rate do play an important role in the overall dynamics, a sufficiently small transcription rate or sufficiently pronounced pausing event after binding is no longer possible gives the system enough time to almost entirely refold to the ON state regardless of the fraction of bound transcripts.

From numerical simulations of continuous transcription in presence of external variations in metabolite concentration, we have observed that the relevant timescale for riboswitch function corresponds to the mean time it takes for the binding window to be traversed, and its relationship to the timescale of changes in metabolite concentration. For slow changes in concentration the system equilibrates at large bound fractions, and the dynamic range of the final fraction ON depends entirely on the transcription rate. This agrees with our intuition and analytical results since, even in total absence of metabolite, if transcription proceeds too quickly the transcripts do not have sufficient time to refold from the OFF to the ON state, so that dynamic range is reduced at a small fraction ON—bottom-right panel of Fig. 4. At the same time, for sufficiently low transcription rates, permitting a large fraction ON in absence of metabolite, fast modulations in concentration result in a reduced dynamic range at a large fraction ON—top-left panel of Fig. 4. A combination of both slow changes in metabolite concentration and low transcription rates, in turn, yields a wide dynamic range—bottom-left panel of Fig. 4. This is not affected by the maximum concentration attained by the metabolite at different values of the stopping time *t*_1_, and using a step function yields the same behavior—see Fig. 5.

**Figure 5:**
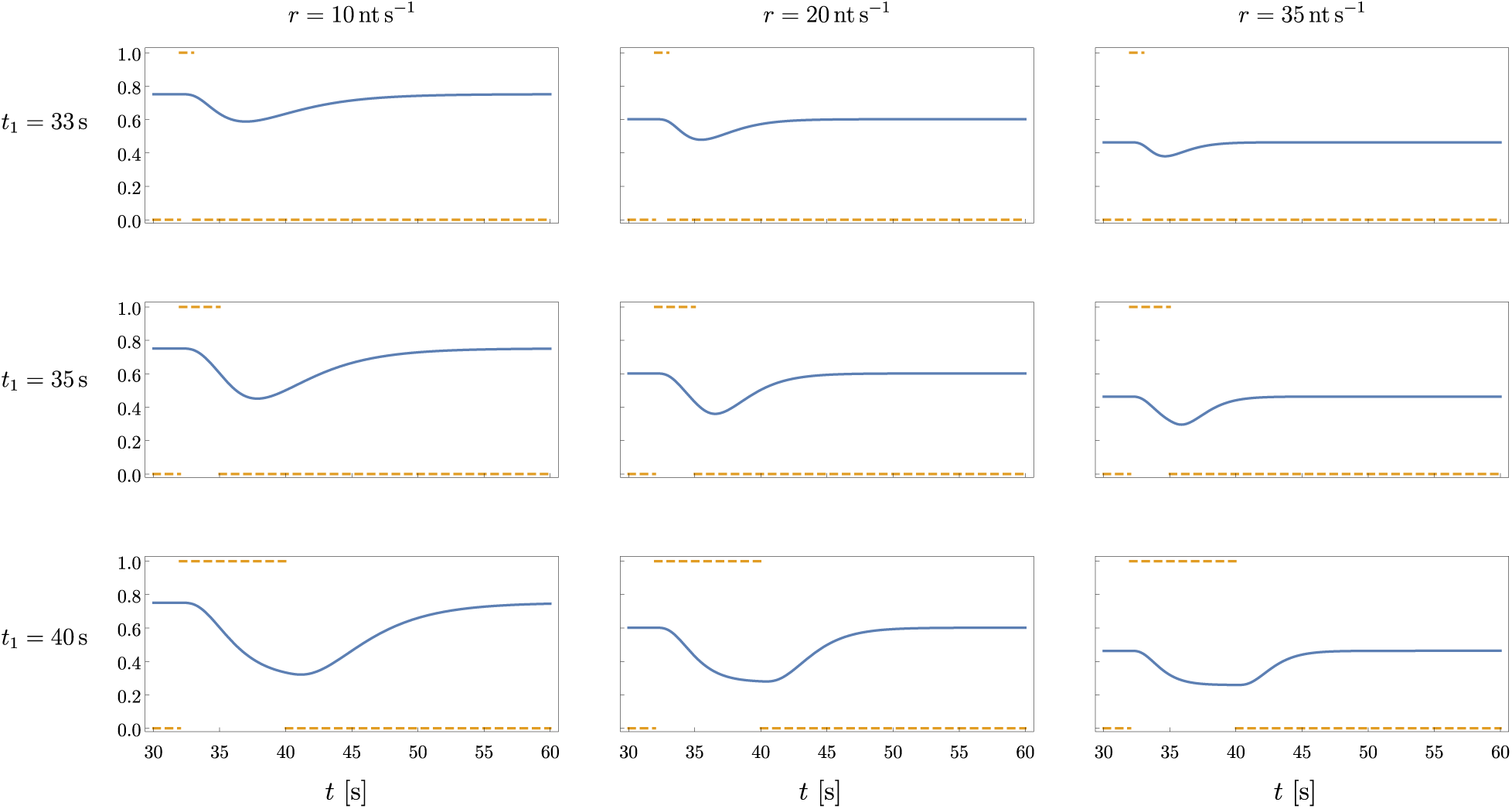
**Orange, dashed line:** the ligand concentration follows a step function, being set to *M* = 1 from a time *t*_0_ = 32 s until a time *t*_1_, and to *M* = 0 otherwise; top: *t*_1_ = 33 s; middle: *t*_1_ = 35 s; bottom: *t*_1_ = 40 s. **Blue, solid line:** resulting fraction ON at a length of 137 nucleotides for the (2’dG)-sensing riboswitch from *Mesoplasma florum* under continuous transcription at a constant rate; left: *r* = 10 nt s^−1^; center: *r* = 20 nt s^−1^; right: *r* = 35 nt s^−1^. An equilibration time of 30 s has been used. The resulting behavior is equivalent to that of Fig. 4. Parameter values: *k*_*b*_ = *µ* = *ν* = 1. Refolding rates from Helmling *et al.* (2018) [8].

While our illustrations have been based on the case of an OFF riboswitch, for which metabolite binding stabilizes the OFF state—thus drastically reducing the folding rate to-wards the ON state—the analytical expressions we have derived are defined in terms of arbitrary values for the folding rates 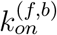 so that, without modifying the available conformations and the allowed transitions between them in the *Mesoplasma florum* example, we can accommodate a simple ON switch that destabilizes the OFF state upon metabolite binding by simply making 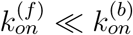. Figure 6 shows the same landscape as Fig. 3A for the case of an ON switch when the values of 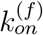 and 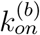 are swapped; Fig. 7, in turn, depicts both cases, OFF and ON, as a function of the metabolite-induced stabilization/destabilization strength—determined by the ratio between 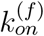 and 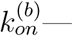for a fixed transcription rate. The different effects of increased metabolite binding—as a result of higher concentration and/or binding rate—for the two cases can be clearly appreciated. In general, the more stable one configuration is with respect to the other, the larger is the dynamic range of the outcome.

**Figure 6:**
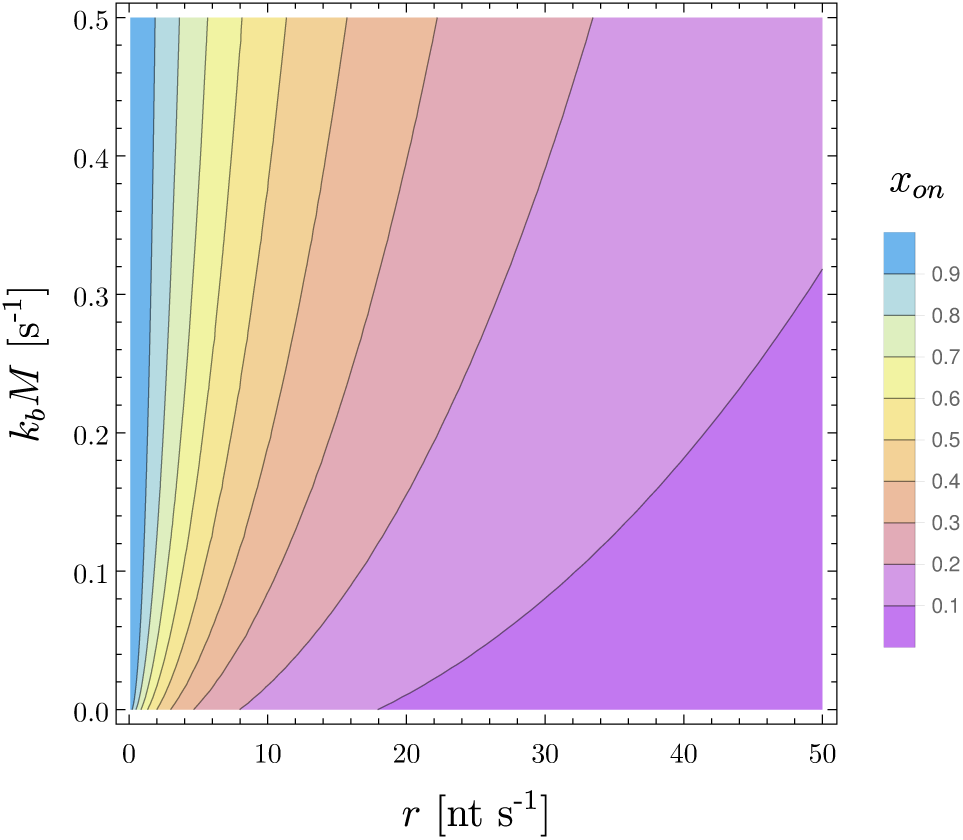
Contour lines for the fraction ON at a length of 137 nucleotides in a single round of transcription for the (2’dG)-sensing riboswitch from *Mesoplasma florum*, as a function of constant transcription rate *r* and the product of the ligand binding rate and the ligand concentration *k*_*b*_ *M*, when the folding rates from the ligand-free and ligand-bound OFF states to the ON state have been swapped, thus resulting in an ON switch. Parameter values from Helmling *et al.* (2018) [8].

**Figure 7:**
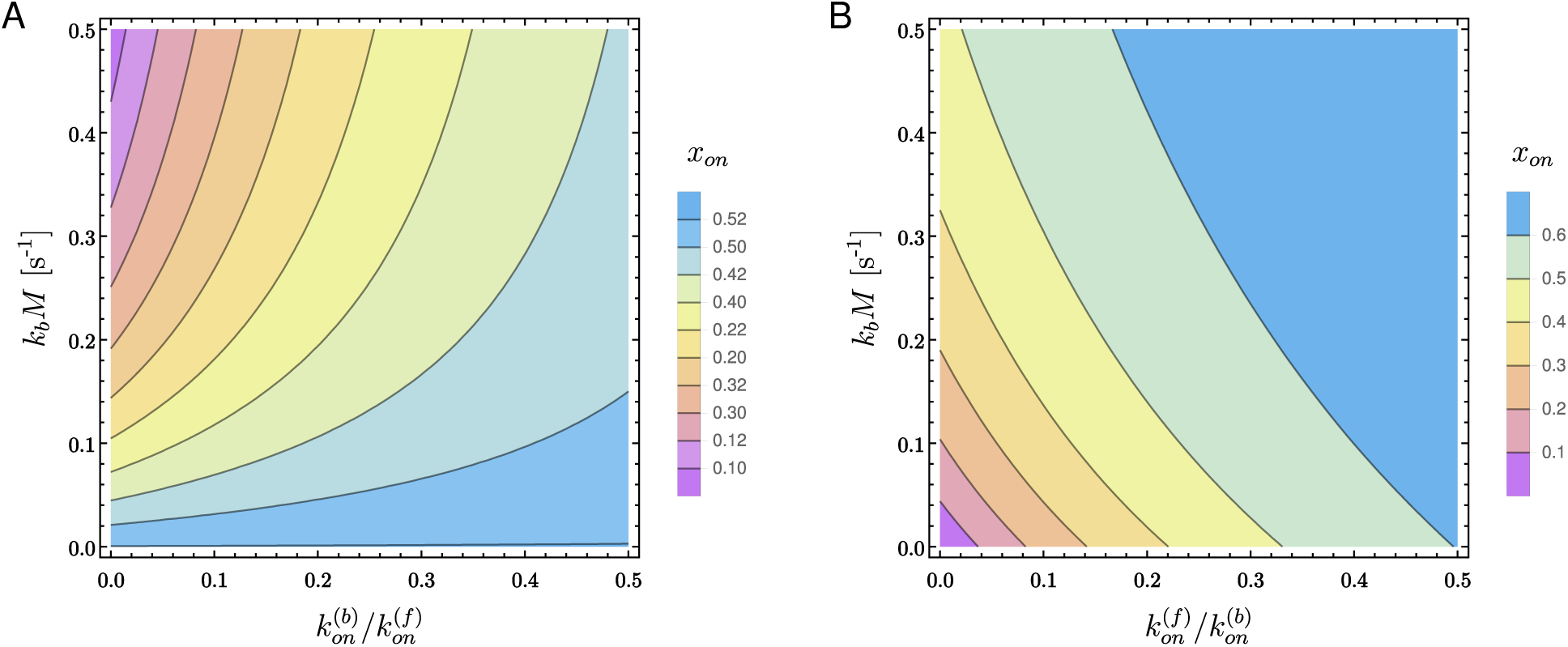
Contour lines for the fraction ON at a length of 137 nucleotides in a single round of transcription for the (2’dG)-sensing riboswitch from *Mesoplasma florum*, as a function of the product of the ligand binding rate and the ligand concentration *k*_*b*_ *M* and the metabolite-induced stabilization/destabilization of the OFF conformation—determined by the ratio between 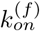 and 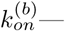for a fixed transcription rate *r* = 10 nt s^−1^. **(A)** Metabolite binding stabilizes the OFF conformation (OFF switch). **(B)** Metabolite binding destabilizes the OFF conformation (ON switch). In both cases, the highest folding rate is set to 1.26 s^−1^.

We have explored the dynamics of the riboswitch under a time-dependent metabolite concentration that is manipulated externally, consisting of a single round of growth, equilibration and decay. While other slower-than-transcription effects—such as temperature dependence of transcription or folding rates [21, 22, 23]—can be directly introduced in the framework of continuous transcription without additional steps, the model we have defined may be sub-sequently coupled to an equation describing the dynamics of metabolite concentration itself as a function of the fraction of transcripts allowing gene expression, in a way akin to that in Santillán & Mackey (2005) [16]. Such a coupling would make possible a full exploration of the feedback mechanisms governing the regulatory behavior of transcriptional riboswitches.

We have also worked under the assumption that the elongation times between transcriptional intermediates are all exponentially distributed, so that the dynamics can be well described by a low-dimensional system of ordinary differential equations (ODEs). The problem may be more complex to model: if we consider single nucleotide addition by the polymerase to be a stochastic process with an exponentially-distributed waiting time, the elongation time from one transcriptional intermediate to another follows a Gamma distribution with a mean equal to the sum of all the individual waiting times [24]; in both cases, transcription proceeds with the same “speed”. We can observe this directly from Eq. 11: even if we choose all transcription rates to be equal, the subdivision of the nucleotide windows into *n* > 1 different sections results in a different form for the fraction ON, compared to that of Eq. 9. For sufficiently large numbers of nucleotides separating all the different transcriptional intermediates, the transitions may be well approximated by a fixed waiting time, as in Hofacker *et al.* (2010) [18]. Within the ODE framework presented here, a full solution may still be obtained recursively from Eq. 9 for the more realistic case comprising windows of 1nt each, traversed in exponentially-distributed times.

## 5. Conclusion

Riboswitches are cis-acting regulatory mRNA elements in bacteria, that modulate the expression of their associated genes in response to a cognate metabolite. They operate either on the level of translation or transcription. Transcriptional riboswitches have to fold into functional structures as they are being synthesized. Here, we developed a mathematical treatment of the kinetic mechanisms of co-transcriptional riboswitch folding. It permits the evaluation of different transcriptional speeds and transcriptional pause sites on population of co-transcriptional folding intermediates and subsequently the functional outcome. The mathematical treatment of the kinetic mechanisms further allows the incorporation of metabolite changes in modelling of the folding pathways. Ultimately, our modelling framework will allow to describe riboswitch based-regulation of transcription at nucleotide resolution.

